# Deep learning of virus infections reveals mechanics of lytic cells

**DOI:** 10.1101/798074

**Authors:** Vardan Andriasyan, Artur Yakimovich, Fanny Georgi, Anthony Petkidis, Robert Witte, Daniel Puntener, Urs F. Greber

**Affiliations:** Department of Molecular Life Sciences, University of Zurich, Winterthurerstrasse 190, 8057 Zurich, Switzerland; MRC Laboratory for Molecular Cell Biology, University College London, Gower St, London WC1E 6BT, UK; Roche Diagnostics International Ltd, Forrenstrasse 2, 6343 Rotkreuz, Switzerland

**Keywords:** artificial intelligence, machine learning, deep learning, predictive infection, computational image analyses, single cell analyses, nucleus, live cell DNA stain, nuclear rupture, nuclear pressure, nuclear envelope, live cell imaging, fluorescence microscopy, laser ablation, adenovirus, herpes simplex virus, virus spreading, virus egress, oncolytic virus, cell lysis, lytic infection

## Abstract

Imaging across scales gives insight into disease mechanisms in organisms, tissues and cells. Yet, rare infection phenotypes, such as virus-induced cell lysis have remained difficult to study. Here, we developed fixed and live cell imaging modalities and a deep learning approach to identify herpesvirus and adenovirus infections in the absence of virus-specific stainings. Procedures comprises staining of infected nuclei with DNA-dyes, fluorescence microscopy, and validation by virus-specific live-cell imaging. Deep learning of multi-round infection phenotypes identified hallmarks of adenovirus-infected cell nuclei. At an accuracy of >95%, the procedure predicts two distinct infection outcomes 20 hours prior to lysis, nonlytic (nonspreading) and lytic (spreading) infections. Phenotypic prediction and live-cell imaging revealed a faster enrichment of GFP-tagged virion proteins in lytic compared to nonlytic infected nuclei, and distinct mechanics of lytic and nonlytic nuclei upon laser-induced ruptures. The results unleash the power of deep learning based prediction in unraveling rare infection phenotypes.

## Introduction

Virus infections give rise to a wide range of cellular phenotypes with complex biology. Depending on the virus, infections can lead to cell cycle and growth enhancement, arrest, cell swelling or shrinkage, membrane blebbing or contraction, and organelle alterations, such the collapse of the secretory pathway, mitochondrial aggregation or nuclear condensation (1–5). Alterations can be pro- or anti-viral, and comprise metabolic changes, such as shifts in glucose, lipid or nucleotide metabolism, as well as apoptotic, necroptotic, and virus-induced cell lysis features (3, 6–8). While programmed cell death can be triggered prior to virus dissemination and has strong anti-viral and pro-inflammatory effects, virus-controlled cell lysis is typically proviral, releases large amounts of progeny virions, and spreads the infection to neighboring cells and between organisms.

Cell lysis has been observed with both enveloped and nonenveloped viruses. It is exploited in clinical applications by oncolytic viruses, such as herpes simplex virus (HSV), measles virus, Newcastle disease virus, vaccinia virus, reovirus, parvovirus, coxsackie virus, polio virus, and adenovirus (AdV) (reviewed in (9)). Yet, virus-induced cell lysis is a short-lasting event, and its detection requires snap shots by ultra-structural analyses and time-lapse fluorescence microscopy with single cell resolution (10–13).

Fluorescence microscopy is a key technology in life science. Many of its original limitations, including sensitivity, sample penetration depth, spatio-temporal resolution and toxicity have been resolved by improvements in optics and chip technology, as well as fluorophore chemistry (14–16). In recent years, it has been further enhanced by artificial intelligence (AI). Particularly deep learning methods allow for complex data analysis, often on par with human capabilities (17, 18). AI and deep learning are increasingly used in biomedical sciences, for example for image recognition, restorations upon low light sampling, and object segmentation(19–25).

Recently, AI applications for pattern recognition in the life sciences have been immensely enhanced by artificial neural networks (ANNs). One of the first implementations of the ANN was the multilayer perceptron (MLP). Inspired by the architecture of the animal visual cortex (26), MLPs are comprised of layers of fully connected nodes (artificial neurons), where each layers output is used as input for succeeding layers. Each neuron of the MLP represents a non-linear function with trainable weights applied to an input signal to generate an output. Regarding image recognition tasks, trained MLPs have been known to perform on par or better than conventional machine learning methods, at the same time being notoriously hard to train and prone to overfitting. The more recent introduction of convolutional neural networks (CNNs) addresses this issue by exploiting the fact that complex patterns in images can be broken to successively simpler features (convolutions). CNNs are composed of three basic building blocks convolutional layers, pooling layers and dense layers. Convolutional layers apply random convolution filters for sub-regions of the input image and derive a feature map, to which a nonlinear function is applied. This layer helps to identify and extract features from the images. The pooling layers down-sample the feature maps extracted by the convolutional layers. This reduces the dimensionality of the data while preserving important information. Dense layers are essentially single layers of an MLP and usually are the final refinement of the CNN, and yield the ultimate classification based on the features extracted from the previous convolutional and pooling layers. CNNs distinguish complex patterns based on reusable network architecture, which makes CNNs versatile for retraining and addressing similar, yet unrelated problems based on new data (27).

One of the current state-of-the-art CNNs uses a residual learning framework to shorten the training time of the networks and make them more accurate. They are called residual neural networks (ResNets) (28). ResNets are widely used by computer vision scientists, and mimic the functionality in the brain, where neurons are arranged in cortical layers and receive input from axonal and dendritic connections in layers distinct from the receiving neuron, thereby skipping other layers. Such skip connection strategies allow a ResNet to further mitigate overfitting and address the problem of vanishing gradient.

Here, we present a ResNet, termed ViResNet, which detects HSV-1 or AdV-infected cells with very high accuracy. Deep learning in ViResNet is qualitatively different from classical machine learning, as it implicitly detects complex morphological features of the infected cell nuclei stained with the DNA-intercalating agent Hoechst 33342, and imaged by fluorescence microscopy (29, 30). HSV-1 and AdV are frequent enveloped and nonenveloped double-stranded DNA viruses, respectively, replicating and assembling progeny capsids in the cell nucleus (31, 32). HSV-1 causes persistent and acute infections of mucosal tissue and a latency phenotype in the trigeminal ganglia of the nervous system (33). Uncontrolled breakout of HSV-1 can give rise to encephalitis, conjunctivitis, Bell’s palsy, eczema, or genital skin lesion. AdVs, in turn, infect the respiratory, ocular and digestive tracts as well as blood cells, and persist in lymphoid cells of human intestines and tonsils (34–36). Upon immunosuppression, they spread to intestinal epithelial cells by unknown pathways, and cause severe morbidity and mortality (37). Our high-fidelity CNN recognizes features of prospective lytic AdV-infected cells, based on Hoechst DNA staining and live cell imaging. The results allowed us to identify physical and morphological features of lytic infected cell nuclei, contrasting nonlytic infected cells. We thereby extend the classical machine learning algorithms, for example, extracting virion trafficking features in cells or particular cell phenotypes (38–43). We provide a resource for analyzing the cell lysis mechanisms in virus infections, a notoriously challenging task owing to the rare occurrence and the inherent terminal phenotype.

## Results

### Machine-labelling identifies HSV-1 and AdV-infected cells in absence of virus-specific staining

To explore the power of deep learning for virus infection biology, we used high content fluorescence imaging and implemented a pipeline scoring the morphology of HSV-1 and AdV-infected cell nuclei stained in live cell mode with the DNA-dye Hoechst 33342 (Fig. 1A, B). We noticed that the Hoechst-stained infected cell nuclei appeared to be distinct from uninfected nuclei. The presence of Hoechst throughout the multi-round infection cycles did not affect the formation of viral plaques (Fig. S1A). We used wild type AdV-C2 and immuno-staining, or the GFP-expressing replication-competent AdV-C2-dE3B-GFP, AdV-C2-dE3B-dADP, AdV-C2-V-GFP, HSV-1-VP16-GFP, HSV-1 C12 viruses (44–46). For AdV, we acquired images at 72 h post infection (pi), and for HSV-1 at 48 h pi, reflecting the different replication times of the viruses. The nuclear images were computationally segmented, and individually annotated according to the infection signal, as AdV or HSV-1-infected or uninfected. They were further processed in a classical image segmentation pipeline yielding individual infected and not infected nuclei (Fig. S1B). This procedure generated up to 2000 cells per condition. We used this dataset to train the computational detection of infected cells using modified architecture cutting-edge network ResNet-50 (28) named ViResNet. We trained two ViResNet classifiers, one for AdV- and one for HSV-1-infected cells (Fig. S1C). To accommodate data of segmented nuclei the input of the network was resized to 224×224 pixels. The ViResNet output layer was adapted to two classes, infected and not infected cells. The hidden layers of the untrained network were initialized using ResNet-50 ImageNet weights. The trained ViResNet had detection accuracies of 96% and 95% for AdV and HSV-1 infected cells, which considerably outperformed the conventional 3-layer multilayer perceptron (3-layer MLP) with accuracies of 81% and 78%, respectively (Fig. 1C). The ViResNet detection classifiers had precision values of 0.94, and recall values of 0.93, respectively, which were much higher than those determined by 3-layer MLP, namly 0.75 and 0.74 for AdV and HSV-1, respectively. The receiver operating characteristic (ROC) curves of ViResNet and the 3-layer multilayer perceptron are shown in Fig. S1D. The trained classifiers were then used on segmented images of nuclei, and thereby enabled the visualization of the infection status at the population level, as well as the quantification of viral spread from infected to uninfected cells, yielding plaque phenotypes (Fig. 1B). In sum, the trained ViResNet accurately identified single AdV and HSV-1 infected cells as well as plaquing cell populations, notably in the absence of specific viral staining, and outperformed the classical 3-layer MLP.

**Fig. 1.**
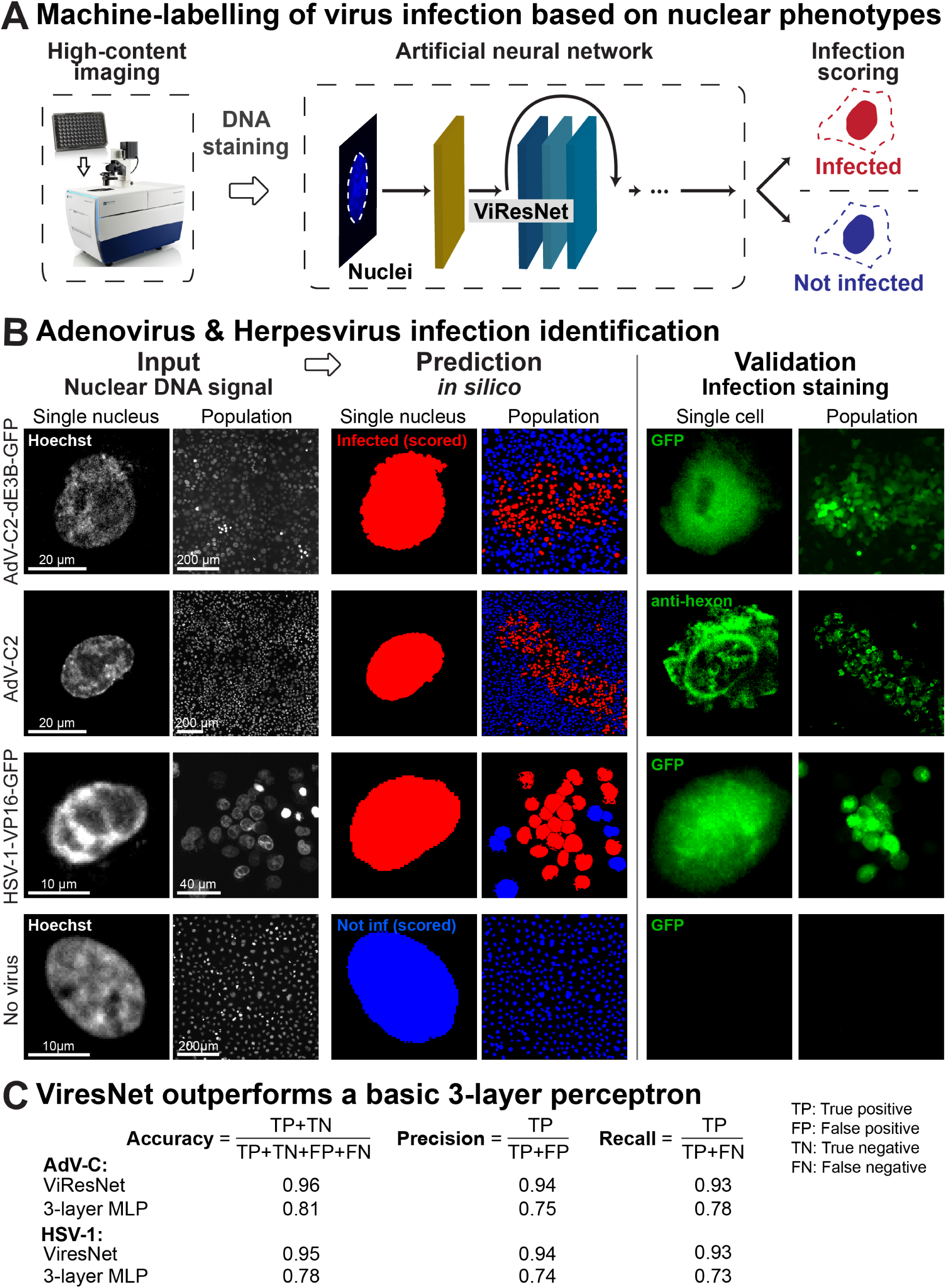
Distinction of HSV-1 and AdV-infected from uninfected cells by machine labelling. (see also Fig. S1). (A) Overview of the machine labelling pipeline for the identification of infected cells. Nuclei stained with Hoechst were acquired using high-content imaging microscope and images fed into ViResnet for classification. (B) Sample of input images, classification results and ground truth data (validation) from GFP transgene expression in case of AdV-C2-dE3B-GFP and HSV-1-VP16-GFP, or immunofluorescence staining for AdV-C2 infection. Uninfected single cells were used as controls. (C) Performance in detection of AdV and HSV-infected cells by ViresNet with regards to accuracy, precision and recall for trained in comparison to a standard 3-layer MLP.

### Lytic egress of AdV from infected cells visualized by fluorescence microscopy

We next employed live cell fluorescence microscopy to analyse the nuclear egress of AdV-C2-GFP-V. These viruses incorporate about 40 GFP-V fusion proteins into their virions, could be purified, and tracked in cells during entry by live cell microscopy (11). Infection of HeLa cells stably expressing histone H2B-mCherry (47) with AdV-C2-GFP-V at multiplicity of infection (MOI) 0.2 showed that newly synthesized GFP-V accumulated in the cell nuclei at about 30 h pi (Fig. 2A, and Movie S1). At 30 to 39 h pi, the localization of GFP-V changed, as indicated by an increased diffuse signal in the cytosol, followed by GFP-V clusters in the cytoplasm. These clusters were largely devoid of H2B-mCherry, which is consistent with the notion that they were made up of AdV particles, which are known to be devoid of histones (11, 48). After the appearance of cytoplasmic GFP-V clusters, the nuclei and the cytoplasm became progressively devoid of GFP-V, and also lost H2B-mCherry, albeit to different extents, indicative of nuclear and cellular lysis, and at least partial nuclear disintegration (Fig. 2A, B, Movies S1 and S2). The quantification of the data showed that about 5.5% of the infected cells developed a lytic nuclear phenotype until 45 h pi (Fig. 2C). The nuclear lysis phenotype was confirmed by EM analyses at 35 h pi revealing clustered AdV-C2-GFP-V particles in the nucleus and the cytoplasm (Fig. 2D). Together the data show that AdV induced nuclear lysis is a rare event and releases clusters of several hundred virions to the cytoplasm and extracellular medium.

**Fig. 2.**
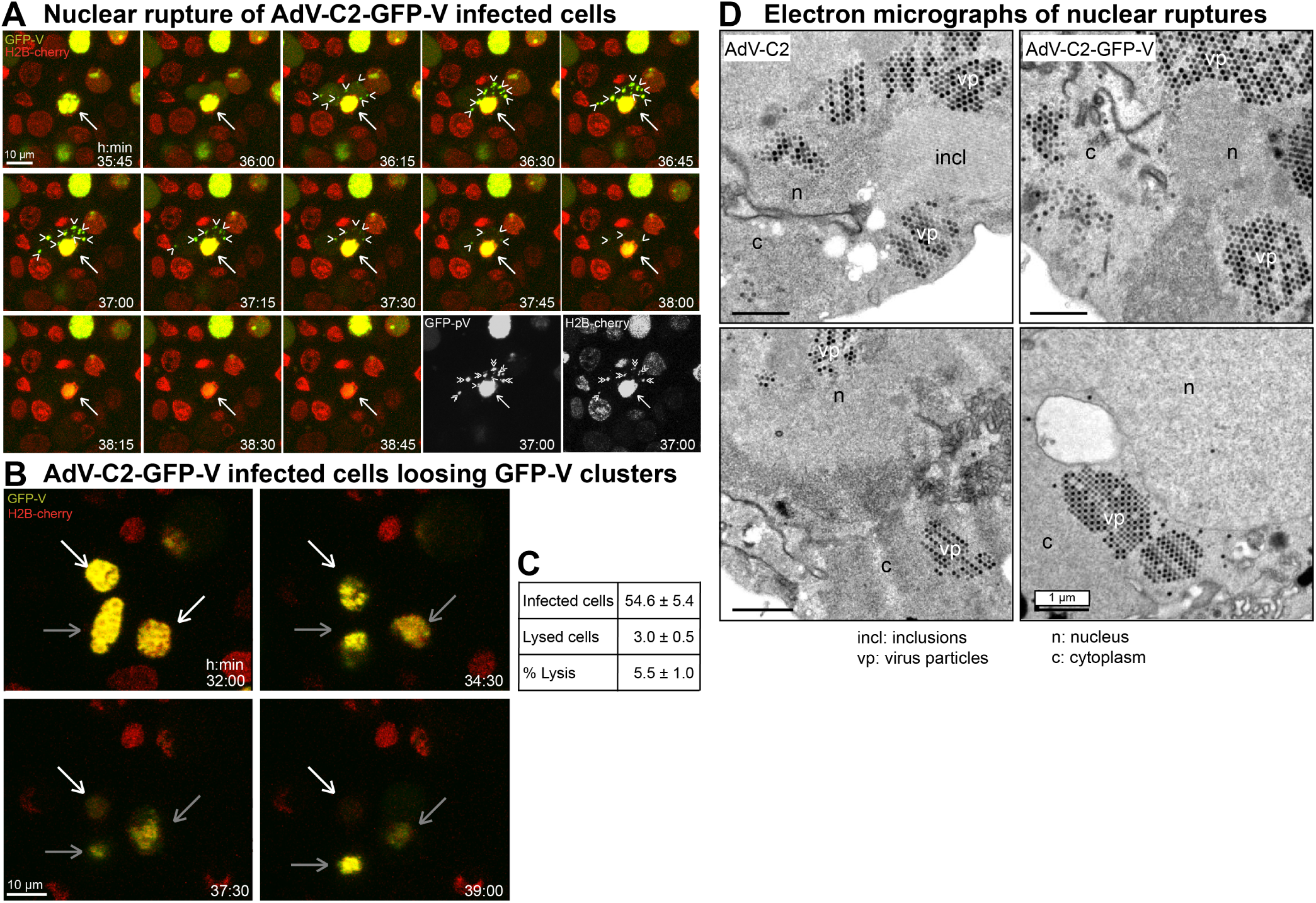
Live cell fluorescence and EM analyses of lytic egress of AdV-C2-GFP-V from HeLa cells. (A) Live cell confocal fluorescence microscopy of H2B-mCherry cells infected with AdV-C2-GFP-V at MOI 0.2 reveals dispersion of GFP-V clusters from the nucleus to the cytoplasm at 35:45 to 38:45 h:min pi. Images show merged maximum projections for GFP-V (green) and H2B-mCherry (red), including the corresponding black/white images at 37 h pi. The white arrow indicates a GFP-V expressing H2B-mCherry positive nucleus undergoing rupture, and arrow heads show GFP-V positive clusters in the cytoplasm. See also Movie S1. (B) Loss of GFP-V clusters from AdV-C2-GFP-V infected cell nuclei. The white arrow points to a GFP-V positive nucleus, which completely looses GFP-V from 32 to 39 h pi. The grey bars denote infected nuclei, which loose GFP-V and H2B-mCherry to a lesser extent in the same time frame. See also Movie S2. (C) Frequency analysis of AdV-C2-GFP-V lytic infected cells. Cell lysis events were scored by the disappearance of both GFP-V and H2B-mCherry signals, quantified from eight different movies at 24 to 45 h pi. (D) Transmission EM images of ultrathin 80 nm sections from epon embedded HER-911 cells 37 h pi. The images show clustered and solitary AdV-C2 (left) or AdV-C2-GFP-V particles (right) in the nucleus (n) and the cytoplasm (c) indicative of nuclear rupture. The virus particles appear in different shades of grey depending on how much of them was present in the section. Also notable are the crystalline-like inclusions (incl) of viral proteins.

### Dataset preparation towards deep learning of lytic and nonlytic infections

Towards harnessing the power of ViResNet to assess and quantify the spreading efficiency of AdV, we employed live cell fluorescence imaging in high throughput mode at very low MOI for up to 4 days. Monolayers of A549-ATCC cells were infected with AdV-C2-dE3B-GFP, AdV-C2-GFP-V, or AdV-C2-dE3B-GFP-dADP, as described (10, 11). The AdV-C2-dE3B-GFP lacks the E3B region encoding the receptor internalization and degradation proteins RID*α* and RID*β* and 14.7K, which are involved in protecting AdV-infected cells against apoptosis and premature lysis by tumor necrosis factor (Gooding et al., 1988; Tollefson et al., 1998). The AdV-C2-dE3B-GFP-dADP additionally lacks the AdV death protein (ADP, E3-11.6K), which is implicated in virus-controlled cell lysis and spreading to neighboring cells (49, 50). All these reporter viruses express GFP from the CMV promoter, replicate and form progeny in cell culture. Accordingly, we monitored the nuclear changes and cell lysis by including Hoechst 33342 and propidium iodide (PI) in the medium, and tracked, segmented and quantified the infected cells using the nuclear Hoechst signal. PI is membrane-impermeable and becomes fluorescent upon binding to nucleic acids in the cytosol or the nucleus of lysed cells (51, 52). The PI stain gave a distinct signal of lysed cells but not of intact cells. As expected (10), the AdV-C2-dE3B-GFP gave rise to comet-shaped plaques 72 h pi, owing to spreader cells, which lysed (Fig. 3A, and also Fig. 2). Remarkably, only about 25% of the infected GFP-positive cells 24 h pi turned out to be spreaders at 72 h pi (Fig. 3A, Fig. 3B). The deletion of ADP reduced the spreader frequency to about 3%, similar as AdV-C2-GFP-V, which expresses GFP-V instead of V and is attenuated in virus propagation (11). Unlike the AdVs, HSV-1-GFP yielded an average of about 85% spreading efficiency, and remarkably this virus gave rise to round-shaped plaques, distinct from the comet-shaped plaques with AdV (Fig. 1B and 3A, 3B). Comet-shaped plaques arise with lytic released but not cell-associated virions carried by convection forces in the culture medium to distant cells (10). We confirmed this notion with the cell lysis indicator PI. A lysing, PI-positive cell always preceded the formation of AdV plaques (Fig. 3C), whereas HSV-1 infected cells giving rise to round plaques had no PI-positive donor cell at their origin (Movie S3), consistent with a nonlytic egress mode of HSV (53). It can be noted that, at later time-points of infection and further rounds of replication (>4 days), additional lytic cells were observed, but it could not be determined if these cells formed plaques, because of the high density of preexisting infected cells.

**Fig. 3.**
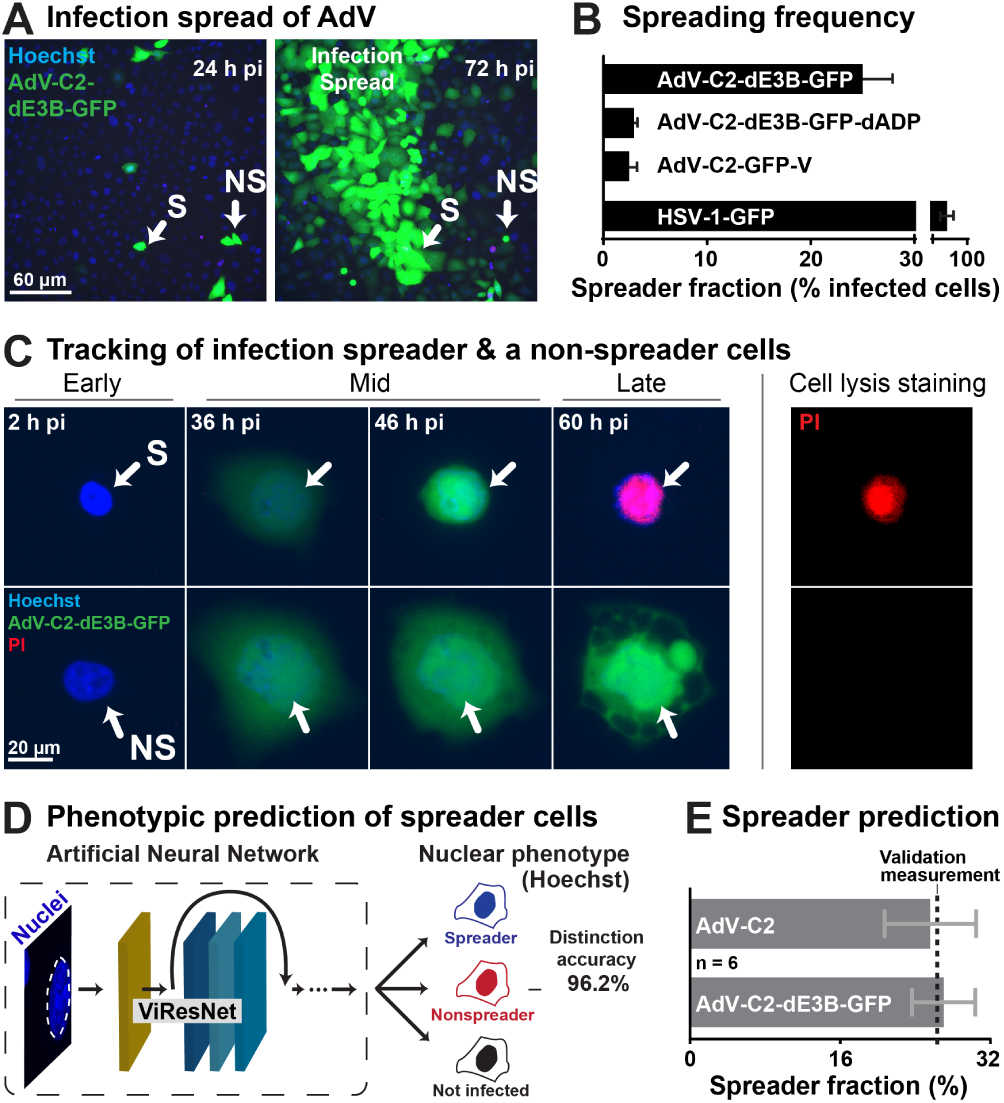
Prediction of AdV lytic and nonlytic infection outcomes based on artificial neural network learning from live cell fluorescence infection analyses. (see also Fig. S3). (A) Fluorescence micrographs displaying the initial infection of human lung epithelial A549 cells (24 h pi) and AdV-C2-dE3B-GFP spread at 72 h pi in presence of Hoechst 33342 staining the nuclei. (B) Measurement of spreader cell fraction for different GFP expressing viruses. Note the reduction of virus-infected A549 spreader cells upon deletion of ADP and in case of AdV-C2-GFP-V, which is known to be attenuated in particle assembly (10). The latter result was in excellent agreement with data from HeLa-H2B-mCherry cells (see Fig. 2C). Note the rather high spreading frequency of HSV-1-GFP infected cells in the absence of lysis of the source cell (PI negative, see Movie S3). (C) Tracking of the infection spreader using live cell fluorescence confocal microscopy of infected A549 cells. Blue indicates nuclear signal (Hoechst), green represents AdV-C2-dE3B-GFP expression and red indicates cell leakiness (propidium iodide, PI). (D) Overview of the processing pipeline in the prediction of spreader and non-spreader cells based on classification of nuclei up to as much as 20 h before lysis. (E) Prediction of spreader fraction in AdV-C2 and AdV-C2-dE3B-GFP infected cells, including the experimentally determined ground truth validation data.

### Deep learning predicts the lytic and nonlytic infection outcomes

Using the live imaging data from three AdV infections, AdV-C2-dE3B-GFP, AdV-C2-GFP-V and AdV-C2-dE3B-GFP-dADP, we constructed a training, validation and testing dataset for a CNN cell fate predictor based on the nuclear Hoechst stain (Fig. S2A). The dataset contained approximately 300 tracked spreader and nonspreader nuclei, each having 10-24 frames of interest (about 4000 images) and additional 2000 images of not infected nuclei. The frames of interest for spreader and nonspreader nuclei were chosen between 24-48 h pi, depending on the onset of morphological changes in the nucleus. We used a modified version of the already established ViResNet to suitably predict the phenotypes. The output layer was changed to three classes - spreading, nonspreading and not infected cells. Feeding test images into the trained network revealed a high level of accuracy (96%), and precision and recall values of 0.95 and 0.952, respectively. This was much higher than the accuracy of the 3-layer multilayer perceptron, as indicated by the ROC curves for identifying the spreader, the nonspreader and the not infected cells among the remaining cells in the population (Fig. S2B). We then utilized the acquired data to construct a pipeline for the prediction of spreader cells for AdV-C2 and AdV-C2-dE3B-GFP infections (Fig 3D). For both viruses the ViResNet predicted the spreader fraction to be about 25% of the infected cells, in excellent agreement with the wet lab data from the AdV-C2-dE3B-GFP infections (Fig. 3E). This result concurred with the notion that the E3B region of the viral genome is not essential for the lytic events (10).

### Faster nuclear accumulation of GFP-V in spreader than nonspreader cells

Having established an AI for the predictive determination of cell-fate, we aimed for an initial characterization of lytic and nonlytic infected cells. Attention of the trained networks may be visualized using class activation maps (CAMs). This can be used to characterize phenotypes learned from different information classes (54). The ViResNet CAMs showed that the Hoechst signal in the nuclear periphery was more pronounced in the nonspreader than the spreader cells, while internal Hoechst stained regions seemed to be similar overall (Fig. 4A). We next analyzed the dynamics of the Hoechst signal and the nuclear size from 0 to 68 h pi, when most of the spreading cells had lysed. While the average Hoechst intensity profiles remained indistinguishable between spreader and nonspreader nuclei, the nuclear area dynamics were distinct, showing a steady area increase for nonspreader nuclei until 30 h pi, followed by a decline until 50 h pi and another increase to 68 h pi (Fig. 4B). In contrast, the area of the spreader nuclei did not significantly increase until 20 h pi, but then rapidly surged till 40 h pi, followed by a steady phase, and a decline a few hours before lysis. Interestingly, the expansion phase of spreader nuclei (annotated by the cell lysis indicator PI) coincided with the rapid accumulation of the viral structural protein GFP-V (Fig. 4C).

**Fig. 4.**
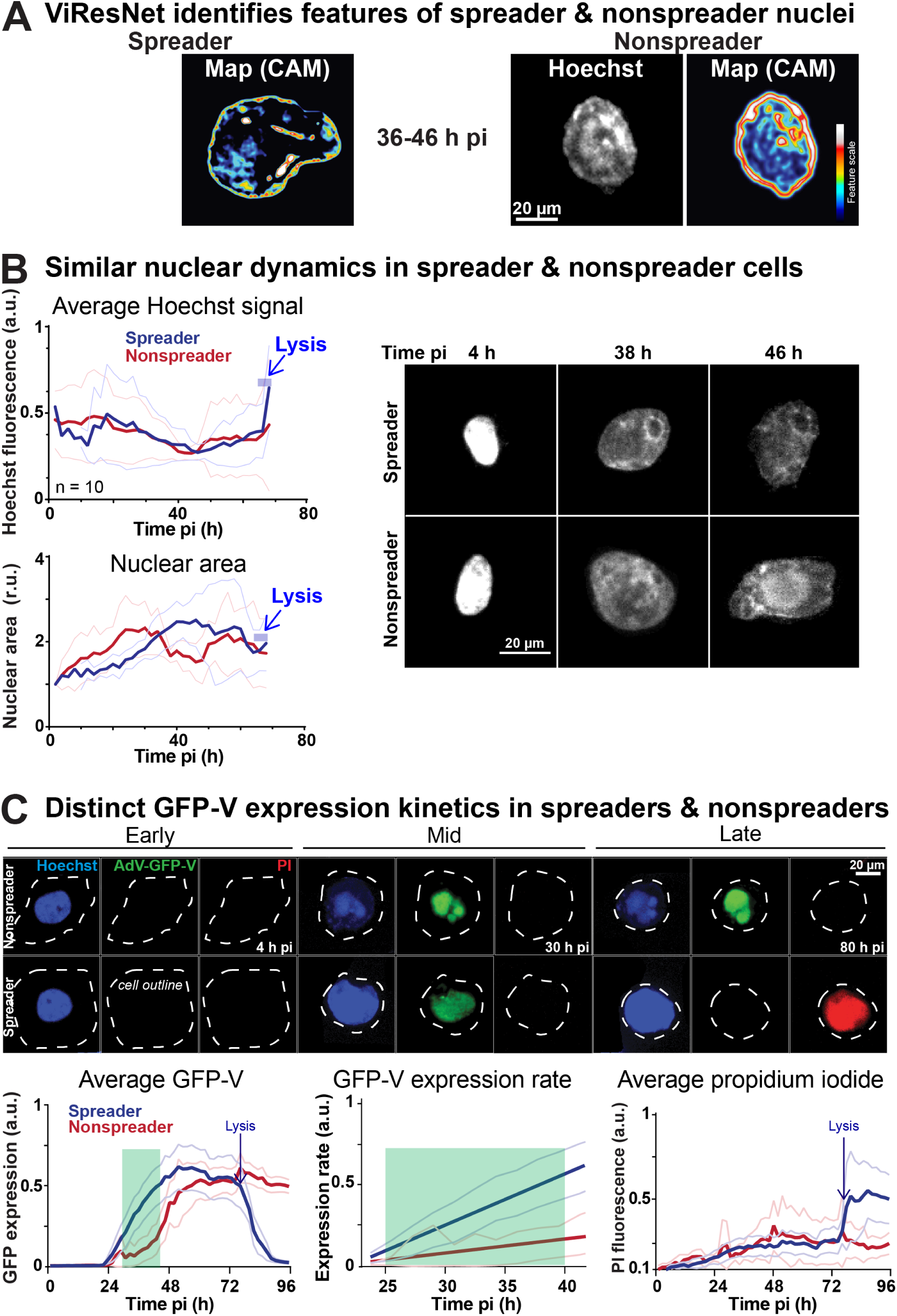
Features of spreader and nonspreader nuclei. (A) The CNN ViResNet defines class activation maps predicting spreader and nonspreader nuclei. (B) Time-resolved analysis of spreader and nonspreader nuclei prior to lysis, based on the average intensity of Hoechst in the segmented nuclear area. Spreader and nonspreader nuclei are notated in blue and red accordingly (n=10). Thin lines denote the standard deviation of the mean. (C) Tracking of AdV-C2-GFP-V infected cells based on nuclear (Hoechst), viral (GFP) and cell lysis (PI) indicators. Thick lines indicate the average normalized intensity of GFP-V signal, or the average normalized intensity of the PI signal (n=30). Thin lines denote the standard deviation of the mean. The expression rate of GFP-V at 24-40 h pi (green area) was derived from linear regression of the average normalized intensity of GFP-V (n=30).

Importantly, GFP-V is a functional fusion protein, and in AdV-C2-GFP-V replaces the wild type V, an inner virion component serving as a linchpin between the virion DNA and the capsid wall, and localizing to the nucleoplasm and nucleolus (11, 36, 55). The data show that spreader and nonspreader nuclei exhibit significant morphological differences, as well as distinct accumulation rates of a viral structural protein GFP-V.

### Spreader cell nuclei have distinct mechanics from nonspreader nuclei

Lytic spreading of AdV entails that the infected cell bursts open and releases virions to the neighboring cells and the environment, as observed in cell culture, animal models and reactivation from persistence in human intestines (10, 37, 56). The lytic release of AdV particles from the nucleus requires the rupture of the nuclear envelope (NE). NE rupture in interphase is commonly defined as the event where the integrity of the nuclear membrane is lost, and nuclear and cytoplasmic proteins rapidly mislocalize, notably in the absence of chromatin condensation. NE typically ruptures in cells with defects in nuclear lamina organization, and under mechanical stress, such as enhanced acto-myosin contractile forces, or in micronuclei, which arise as a consequence of mitotic pathology. The latter is often caused by unrepaired DNA damage, chromatin mis-segregation or breakage of chromatin bridges at the end of mitosis (57, 58). NE rupture is also thought to release intranuclear pressure when cells migrate through small constrictions, and facilitate nuclear deformation (59). To assess mechanical features in spreader and nonspreader infected nuclei, we experimentally breached the integrity of the NE by laser ablation microscopy of AdV-C2-GFP-V infected cells in presence of the nuclear dye Hoechst (Fig. 5A, and Fig. S3). Prior to ablation at 40 h pi, live cells were classified on-the-fly as either spreaders or nonspreaders using the above developed CNN. This was then followed by a focused illumination with an intense laser beam (laser ablation) at invariant intensity and size bursting open a defined area of the nuclear envelope The volume of the nuclei and the exerted nuclear contents indicated by the cytoplasmic area of Hoechst-positive material were recorded by fluorescence imaging at low intensity in live cells. Results indicated that rapidly after ablation (<1min), the prospective lytic nuclei lost more than 50% of their original volume, whereas the nonlytic or uninfected nuclei collapsed only minimally (Fig. 5B). This suggested that the mechanical strength of the NE was lower in lytic than nonlytic infected nuclei. To estimate if the relative nuclear pressure across the NE was higher in the lytic than nonlytic infected cells, we measured the area of exerted Hoechst-positive material in the cytoplasm upon laser rupture of the NE. The spreader cells were found to extrude Hoechst-positive material amounting to about three-fold the extrusion area in the nonspreader cells, whereas uninfected cells released only a minimal amount of Hoechst-positive material to the cytoplasm upon ablation (Fig. 5C). This suggested that the NE experienced higher macromolecular pressure from the nuclear contents than for the cytoplasmic side. The results show that the nuclei of lytic infected cells have mechanical properties distinct from the nonlytic cells. These differences are the result of lower NE stability and higher nuclear to cytoplasmic pressure ratio than nonlytic cells. This raises the possibility that the changes in the mechanical properties of nuclei and possibly the cytoplasm are an important determinant for effective cell lysis, and rapid spread of infection.

**Fig. 5.**
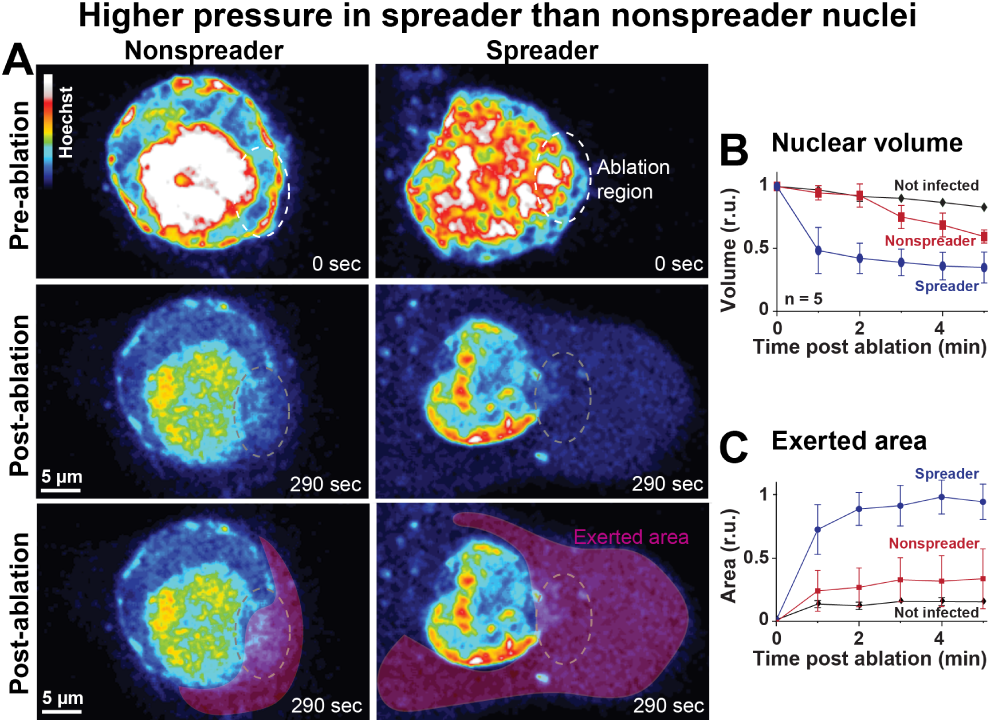
Modulation of nuclear mechanics by AdV. (see also Fig. S5). (A) Live cell imaging of pseudo-colored Hoechst nuclear signal in nuclei identified as spreader and nonspreader, 5 min before laser ablation of the nuclear envelope, as well as 290 sec after the laser ablation, denoted as the ablation region. The lower row highlights the Hoechst-positive material extruded into the cytoplasm upon laser ablation, and is highlighted by the extruded area. (B) The normalized nuclear volume was measured using Hoechst signal every minute post ablation up to 5 min by normalizing the segmented volume to the initial volume (n=5). (C) The exerted area was measured as exerted background area of the Hoechst signal post ablation normalized by the segmented initial area of the z-stack maximum projection of the nucleus (n=5). Error bars represent standard deviations. Spreader, nonspreader and not infected nuclei are denoted in blue, red and black, respectively.

## Discussion

The dynamics of viral genomes in cells are virus family-specific and essential for infection. Retroviruses, such as the human immunodeficiency virus, or negative sense RNA-viruses, such as influenza virus use the nuclear export machineries of the cell to transport their replicated genomes from the nucleus to the cytoplasm in continuous processes (60, 61). Viral genomes in a nucleocapsid, such as AdVs, parvoviruses, polyomaviruses exit from the nucleus by rupturing the NE (49, 62–65). Others, for example herpesviruses and baculoviruses, egress from the nucleus by engaging distinct budding processes (reviewed in (15, 66–68)).

Yet, in the absence of the nonlytic nuclear egress complex, herpesviruses have been reported to engage processes of nuclear rupture or dilation of nuclear pores (12, 69–71). The underlying mechanisms have been difficult to explore, largely because of cell-to-cell variability, unpredictable onset of lysis and a magnitude of pro- and anti-viral factors precluding the lytic phenotypes in population assays.

### Convolutional neural networks in image recognition and analyses

We focused on single cell analyses, and developed CNNs to identify virus-infected cells, and in a second step identified and characterized lytic infected cells. CNNs have been widely employed in computer vision and image recognition with unprecedented accuracy (72). Beyond merely separating signal from noise in a linear fashion, these CNN algorithms autonomously detect complex patterns in the data from high level user annotations (18). Deep learning algorithms iterate thousands of samples with remarkable robustness and accurately annotate previously hidden features beyond human analytical capability, with recent examples extending to biological and clinical settings (73–75).

### ViResNet, a machine-based procedure identifies virus-infected cells

We developed CNNs for virus infections, and named them ViResNets. They identify distinct phenotypes in the nuclei of cells infected with either HSV-1 or AdV. Both agents replicate and form progeny nucleocapsids in the interphase nucleus, and affect the host chromatin in a virus-specific manner, for example by enriching heterochromatin in the nuclear periphery of the infected cells (65, 76, 77). A similar phenomenon has also been observed with subnuclear structures in polyomavirus infected cells (78). We conjecture that an adapted ViResNet will be applicable to other viruses, beyond AdV and HSV-1. Using the live cell nuclear dye Hoechst 33342 and fluorescence imaging, the procedure unequivocally identified AdV and HSV-1 infected cells, notably in the absence of virus-specific labels. Hoechst 33342 is a versatile bisbenzimide fluorescent compound, commonly used in flow cytometry and fluorescence microscopy, for example to identify S-phase cells with duplicated genome contents. Bisbenzimides stain nuclear double-stranded and cytoplasmic organellar DNA by hydrophobic interactions with adenosine-thymidine rich regions (79, 80). While Hoechst 33342 inhibits vaccinia virus infection, it does not affect HSV-1 or AdV infections (10). Our results open opportunities for diagnosis of viral infection phenotypes in primary tissue beyond cell cultures, and show that Hoechst 33342 is well suited for live cell infection studies addressing the lytic and nonlytic infection phenotypes of AdV.

### Visualizing the lytic nuclear egress of AdV, a non-isotropic process

Using live cell microscopy and electron microscopy, we found that a small fraction of cells released large clusters of AdV-C2-GFP-V particles from the nucleus at late stages of infection, eventually exiting the cell upon disruption of the plasma membrane. This suggests that lytic virion release is a nonisotropic process involving the rupturing of the NE. The lytic nature of AdV egress was corroborated by live cell imaging showing that each AdV plaque within about 4 days of infection emerged from an infected cell that turned PI-positive just a few hours before the onset of plaquing. EM analyses revealed large discontinuities of the NE, and coincidentally cytosolic and also nuclear virion clusters, and crystalline-like inclusions of virion-free viral proteins, in agreement with earlier reports (81). Clusters and inclusions were observed in both AdV-C2 and AdV-C2-GFP-V infections. Cytosolic clusters were micrometers in size, and resembled the large fluorescent AdV-C2-GFP-V puncta, which were short-lived for 1-2 hours before they dispersed from the cells. The clusters had the appearance of densely packed pseudocrystals comprising some hundreds to thousands of virions. We speculate that the pseudocrystals are formed by phase separation in the nucleus in a coarsening process of interfacial area reduction, known as ‘Ostwald ripening’, according to the physical chemist W. Ostwald who first described the process (82). ‘Ostwald ripening’ involves the minimization of total interfacial energy between components, and drives crystal growth from precipitate clusters in inorganic systems and organic crystals, such as tobacco mosaic virus (83, 84). The AdV pseudocrystals disappeared upon lytic release from cells, and rapidly dispersed in the extracellular medium. Cells in the vicinity of a lytic infected cell are thus likely to be exposed to a high multiplicity of virus particles, a conjecture which is in agreement with the observation that these cells rapidly express viral GFP from a transgenic promoter within a couple of hours after lysis of the founder cell.

### Convolutional neural networks identify distinct mechanical features of lytic infected cells

Unequivocal assessment of infection phenotypes and molecular analyses are drivers for biomedical discovery and disease management. Specific phenotypic reporting of infections typically occurs by immunolabeling reagents in diagnostics, or by transgenic viruses expressing a fluorescent of luminescent protein in research. To enable studies of infected cells undergoing lysis, we developed a combination of low-invasive high-throughput confocal microscopy and CNNs based on Hoechst-stained nuclei in time-lapse series of infected cells. These procedures allow the prospective identification of lytic infected cells and distinct features of lytic cells. The lytic release of AdV from the nucleus involves ADP, and possibly limited proteolysis of NE proteins (50, 85–87). ADP is a small protein with a single transmembrane segment and cytosolic C-terminal domain (50). It is critical for AdV-C2/5 lytic infection, nuclear lysis and the production of large viral plaques (88). Moreover, ADP overexpression from infectious viral genomes leads to enlarged plaques and enhanced cytopathic effects (87), although the exact mechanism of ADP promoting lytic infection is unknown. We show that lytic infected cells enriched the viral structural protein GFP-V in the nucleus at faster rates than nonlytic infected cells. The levels of GFP-V in lytic and nonlytic cells reached similar levels, and in AdV-C2-dE3B-GFP cells GFP accumulated to similar extents in lytic and nonlytic cells, suggesting that viral protein expression alone is not a discriminator of the lytic/nonlytic infection outcome. Our laser rupture experiments of the NE, however, indicated that the lytic infected nuclei shrunk to greater extent and expelled their chromatin contents further to the cytoplasm than the nonlytic cells. The former implies that the nuclear envelope including the chromatin in the nuclear periphery of lytic cells is less stable than in nonlytic cells. It can now be tested if the peripheral heterochromatin is preferentially depleted in lytic cells, or if the LINC (linker of nucleoskeleton and cytoskeleton) complex in the NE connecting the cytoplasmic and the nuclear filaments is impaired (89, 90). In addition, the laser ablations showed that the difference between the macromolecular pressure inside the nucleus and the cytoplasm is larger in the lytic cells than the nonlytic cells. We speculate that proteins and nucleic acid polymers in the nucleus exert osmotic pressure, which facilitates the expulsion of the virus clusters, possibly enhanced by host chromatin fragmentation and decompaction, as well as redistribution of proteins of the LINC complexes in the NE (91–93). It is now feasible to dissect the more than one hundred known protein interactions between lamins and the inner nuclear membrane, most of which are integral membrane proteins harbouring LEM and SUN domains (94, 95). The virus clusters released from the nucleus frequently remained associated with the cytoplasm, before they displaced to the medium, which suggests the disruption of the plasma membrane after nuclear rupture. Nuclear rupture may release the viral cysteine protease L3/p23 to the cytosol, and thereby enhance degradation of the cytoskeleton (96). L3/p23 cleaves cytokeratin K18, microtubules and actin, and makes the infected cell more susceptible to mechanical stress (97). Although cellular lysis is required for AdV spread in tissue, it occurs only in a small fraction of initially infected cells over a given time frame (10). Precisely how the plasma membrane is disrupted and the virus clusters released from the cell can now be analysed in detail. In summary, our deep learning procedure is versatile and in principle adaptable to any kind of pathogen infection and can be combined with single cell analysis technologies. It allows for the targeted interference with and analyses of cryptic infection phenotypes, and ultimately the discovery of thus far hidden infection phenotypes.

## ACKNOWLEDGEMENTS

We thank Silvio Hemmi for generating AdV-C2-dE3B-GFP-dADP, Ivo Sbalzarini and Thomas Greber for fruitful discussions. Nicole Meili, Melanie Grove, Corinne Wilhelm and Karin Boucke are acknowledged for expert help with tissue culture and electron microscopy. The work was supported by an excellence grant from the Swiss National Science Foundation to UFG (31003A_179256 /1), the Sinergia project from the Swiss National Science Foundation to UFG (Paracrine delivery of therapeutic biologicals, CRSII5_170929 /1), and the Kanton of Zurich.

## Materials and Methods

### Cell culture

A549-ATCC cells were cultivated in a T75 flask and incubated at 37°C, 5% CO2 and 95% humidity (standard conditions) in DMEM medium supplemented with 8% FCS, 1% P/S and 1% L-glutamine. The culture was passaged by splitting 1:10. The old medium was removed, cells were washed with PBS and briefly soaked in 1.5 ml trypsin (Sigma-Aldrich) and incubated at 37^*◦*^C for 3 min. Trypsin was inactivated by adding fresh medium, the cells resuspended and seeded in fresh plates. Hela cells stably expressing histone H2B-mCherry were kindly obtained from Daniel Gerlich (47).

### Viruses

AdV-C2-dE3B-GFP, AdV-C2-GFP-V and AdV-C5-dE1-GFP were grown as described (10, 11, 98). HSV-1-GFP was generated by S. Efstathiou (University of Cambridge, Cambridge, United Kingdom) and kindly provided by C. Fraefel (University of Zurich, Zurich, Switzerland). AdV-C2-dE3B-dADP was generated using two recombineering steps (10, 99, 100). At first, the GalK cassette was introduced into pKSB2-AdV-C2-dE3B-GFP to replace the ADP sequence. The GalK cassette was amplified using the forward primer 5’-ACTGCAAATTTGATCAAACCCAGCTTCAGCTTGCCT-GCTCCAGAGcctgttgacaattaatcatcggca-3’ and the reverse primer 5’-GAACTAATGACCCCGTAATTGATTAC-TATTAATAACTAGTCTCATctcagcactgtcctgctcctt −3’ introducing 45 nucleotides of flanking sequences. Subsequently, the GalK sequence was replaced with a dsDNA of the sequence actgcaaatttgatcaaacccagcttcagcttgcctgctccagagatgagactagttattaatagtaatcaattacggggtcattagttc resulting in deletion of ADP. To generate infectious virus, circular pKSB2-AdV-C2E3BGFPdADP was transfected in human 911 cells stably expressing I-SceI endonuclease (101) using the JetPEI transfection reagent (Polyplus transfection, Illkirch-Graffenstaden, France). Constitutive I-SceI expression in these cells was accomplished following transduction with MLV-ER-I-SceI-HA, which encodes a form of the endonuclease that can be translocated to the nucleus upon treatment with 4-OH-tamoxifen 3h post transfection (102). Cells were selected in medium containing puromycin at 1µg/ml and bulk cultures were expanded under selection conditions. I-SceI expression was confirmed by Western blotting of whole cell lysates using the anti-HA antibody (HA.11 clone 16B12, Covance). Resulting recombinant AdV-C2-dE3B-dADP was plaque-purified and amplified, followed by two rounds of CsCl purification (103). Loss of ADP expression was confirmed by Western immunostaining using the rabbit anti-ADP 87-101 antibody, kindly supplied by Dr. William Wold [Saint-Louis University, Saint-Louis, USA, (87)].

#### EM analyses

HeLa cells grown on alcian blue coated coverslips at 90% confluency were infected with AdV-C2 or AdV-C2-GFP-V at moi 5 for 40 h. Cells were fixed in glutaraldehyde, and prepared for Epon embedding and EM analyses as described (104).

#### Quantification of spreader ratio

To quantify the plaque formation efficacy, the number of infected cells after the first round of infection and the number of plaques developed in the subsequent infection cycles were assessed. AdVs and HSV were titrated on A549-ATCC cells. Monolayers were seeded in 96-well plates (at 15,000) in 100 µl medium overnight. The next day, virus stocks were diluted 1:10,000, followed by a 1:2 serial dilution into medium. The cell supernatant was removed and 100 µl virus dilution or medium added to the cell wells. Following 30 min of incubation at 37°C, the supernatant was replaced by 100 µl fresh medium for endpoint measurements (AdV-C2-GFP-V) or fresh live cell imaging medium (phenol red-free DMEM medium supplemented with 8% FCS, 1% P/S, 1% L-glutamine, 250 ng/ ml Hoechst 33342 obtained from Sigma and 3 µM DRAQ7, the latter obtained from Abcam). The cells were incubated at standard conditions. For endpoint immunofluorescent quantification of AdV-C2-GFP-V, two separate plates were infected and fixed in 4% PFA supplemented with 6 µg/ml Hoechst 33342 at the indicated timepoints, followed by staining with the mouse anti-AdV-C2 hexon antibody (MAB8052, Sigma-Adrich) and anti-mouse Alexa594 (ab150116, Abcam) prior to imaging. Imaging at the indicated time-points was performed with a Molecular Devices IXM-XL/C microscope, using a 4x objective. The number of infected cells and the number of plaques in a well were scored using Plaque 2.0 software (105). The detection of infected cells was verified using CellProfiler software (106). Virus dilutions were chosen such, that well emerging plaques were well separated and could be unequivocally assessed. The following timepoints for first and second replication cycles were used: AdV-C2-dE3B-GFP 48 and 76 h post infection (pi), AdV-C2-GFP-V 48 and 78 h pi, AdV-C2-dE3B-GFP-dADP, and AdV-C5-dE1-GFP 48 and 72 h pi, HSV-1-GFP 14 and 27 h pi, respectively. Spreader ratios were calculated for each well by dividing the plaque number (second replication cycle) by the number of infected cells at first round of replication multiplied by 100.

#### Live-cell microscopy of AdV-C2-GFP-V

Live-cell laser scanning fluorescence microscopy was conducted with an inverted Leica SP5 confocal microscope (Leica Microsystems, Switzerland) equipped with a 63x (oil immersion, NA 1.4) objective, a diode laser (405 nm excitation), an argon laser (458/476/488/496/514 nm excitation), a helium laser (561/594/633 nm excitation) and a humidified chamber at 37^*◦*^ C and 5% CO_2_. HeLa cells stably expressing H2B-mCherry (47) were plated at 25% confluency onto 35 mm glass bottom microwell dishes, and cells allowed to adhere overnight, followed by infection with AdV-C2-GFP-V (0.8 µg) for 24 h. Image stacks were recorded every 15 min for 21.5 h and processed with ImageJ. From 8 movies cell lysis events were determined and plotted against total numbers of infected cells as judged by the expression of GFP-V.

#### Time-lapse live cell imaging for machine learning infection phenotypes

A549 human lung carcinoma cells where seeded into a 96-well plate to achieve 70% confluency. At about 24 h post seeding cells were inoculated with virus for 30 min, virus washed off and cells replenished with fresh phenol red-free DMEM supplemented with nuclear dyes Hoechst (250 ng/ml) and propidium iodide (100 ng/ml). The cells were monitored every 1.75 h for 4 days using a Molecular Devices IXM-C high-throughput confocal microscope. Images were acquired using a 20x air objective. In each well 36 sites were acquired to cover 85% of the well area. At each time-point, one image was generated using a maximum projection of 8 slices of the sample spanning 30 µm along the Z-axis. To identify and classify the features characteristic for AdV lytic and non-lytic infection, a time-lapse experiment was performed in human lung carcinoma A549 cells. Cells were inoculated with AdV-C2-GFP-V, washed, and imaged in presence of the nuclear dye Hoechst and the cell lysis marker propidium iodide, as described earlier (10, 11, 105), except that this time a less invasive high throughput spinning disc confocal microscopy protocol was used (Molecular Devices IXM-C). The cells were monitored every 1-2 hours for 4 days, and images acquired using a 20x air objective. At each time point, an image was computed using a maximum projection of 8 optical slices spanning across 30 µm along the z axis. The projected images were loaded into Fiji image processing environment (107), and infected nuclei centroids tracked utilizing the Trackmate plugin (108). The derived tracks were curated either as spreading or non-spreading depending on presence or absence of the propidium iodide staining at later time points of infection. As the tracks only contain information about the centroid of the nuclei and not the entire segmented nuclei, additional segmentation was performed using custom segmentation pipeline implemented in python (https://github.com/viresnet/). The analysis of the segmented data revealed that late in infection (40 h pi) nuclei will showed biggest morphological differences between spreading and non-spreading cells for both virus and nuclear signals. Therefore, images for this timeframe were used to perform training, validation and testing of the CNN.

#### Quantification and tracking of live infected cells

The acquired image projections were loaded into the Fiji image processing environment (107), and infected nuclei centroids were tracked utilizing the Trackmate plugin (108) and Python-based Trackpy library (https://soft-matter.github.io/trackpy/v0.3.2/). The derived tracks were labelled either as spreading or non-spreading, depending on presence or absence of the propidium iodide staining. As the tracks only contain information about the centroid of the nuclei and not the entire segmented nuclei, additional segmentation was performed using customized Python segmentation functionality. Using the acquired segmentation mask, the average intensities and standard deviations were calculated.

#### Convolutional neural network and training dataset preparation

Based on published the state of the art convolutional neural network for image recognition (ResNet-50) (28) we developed ViResNet using the Python programming language high-level deep learning library Keras (https://keras.io/) employed on top of Tensorflow (see (109)). The architecture was modified by changing the last fully-connected layer succeeding global average pooling layer to either 2 or 3 dimensions, depending if the network was intended to use to classify infected/not infected nuclei or predicting spreading/non-spreading/not infecting nuclei. The training sets acquired by fluorescence microscopy were augmented and expanded (random rotations, shifts, flips) using tools provided by the Keras library. Training set images for infected nuclei classification ViResNets for HSV-1 and AdV-C2, were comprised of AdV-dE3B-GFP, AdV-dE3B-GFP-dADP, AdV-C2-GFP-V, AdV-C2 and HSV-1-C12/HSV-VP16 infected nuclei and uninfected nuclei stained Hoechst. Images were acquired at 48 and 72 h pi for AdV-C2 and 24 and 48 h pi for HSV-1. Training set images for predicting spreader and non-spreader phenotypes using ViResNet were comprised of AdV-dE3B-GFP, AdV-dE3B-GFP-dADP and AdV-C2-GFP-V. Infected cell nuclei with known fate were chosen at 16-36 h pi for AdV-dE3B-GFP, and 28-48 h pi for AdV-C2-GFP-V. To provide sufficient data for the training we expanded the datasets using random augmentations making it to sum up to 40’000 images in total. The resulting dataset was split into 30’000 training, 8000 validation and 2000 test images. All images were downscaled to 224×224 and converted to 3-channel RGB 8-bit in order to correspond to ViResNet architecture input layer. The grayscale to 3-channel conversion was done by duplicating the original grayscale image into red,green and blue channels. During the training ViResNet was initialized using the pretrained ResNet-50 weights for ImageNet competition and Adam optimizer was employed using the default values (110). Training was done using a batch size of 32 images and was stopped as soon as the accuracy and loss reached saturation (1000 epochs). The source code is available at https://github.com/viresnet/.

### Laser ablation of infected cells and quantification of volume change after ablation

A549 cells were seeded on a glass bottom 35 mm petri dish, infected with AdV-C2-GFP-V, and stained with HOECHST dye about 24 h pi. The imaging and laser ablation was performed on a Leica TCS SP8 Multiphoton microscope using Insight DS+ Dual (680-1300 nm and 1041 nm) ultrafast NIR laser tuned to 800 nm and operating at 1% of maximal power of 1.3 W. Ablation was conducted at 20% of the maximal power at 36 h pi. Prior to ablation a Z-stack was acquired in both nuclear and viral channels, which was then fed into a custom Python script which converted it into a maximum projection and fed it to the pre-trained CNN to determine the type of infected nucleus, that is lytic or non-lytic. After the ablation, the nuclei were imaged every minute up to 5 min in the Hoechst channel. The acquired Z-stack images were segmented into nuclear and exerted material volumes using Arivis and Fiji software packages (107).

## Appendix 1 Supplementary Information

This is a supplementary section to the preprint manuscript by Andriasyan & Yakimovich et al. Supplementary Videos can be downloaded alongside with the manuscript. Source code used in this manuscript can be obtained from https://github.com/viresnet/. Supplementary movies are available under https://viresnet.github.io/movies. All further information is available upon request.

**Sup. Fig. 1:**
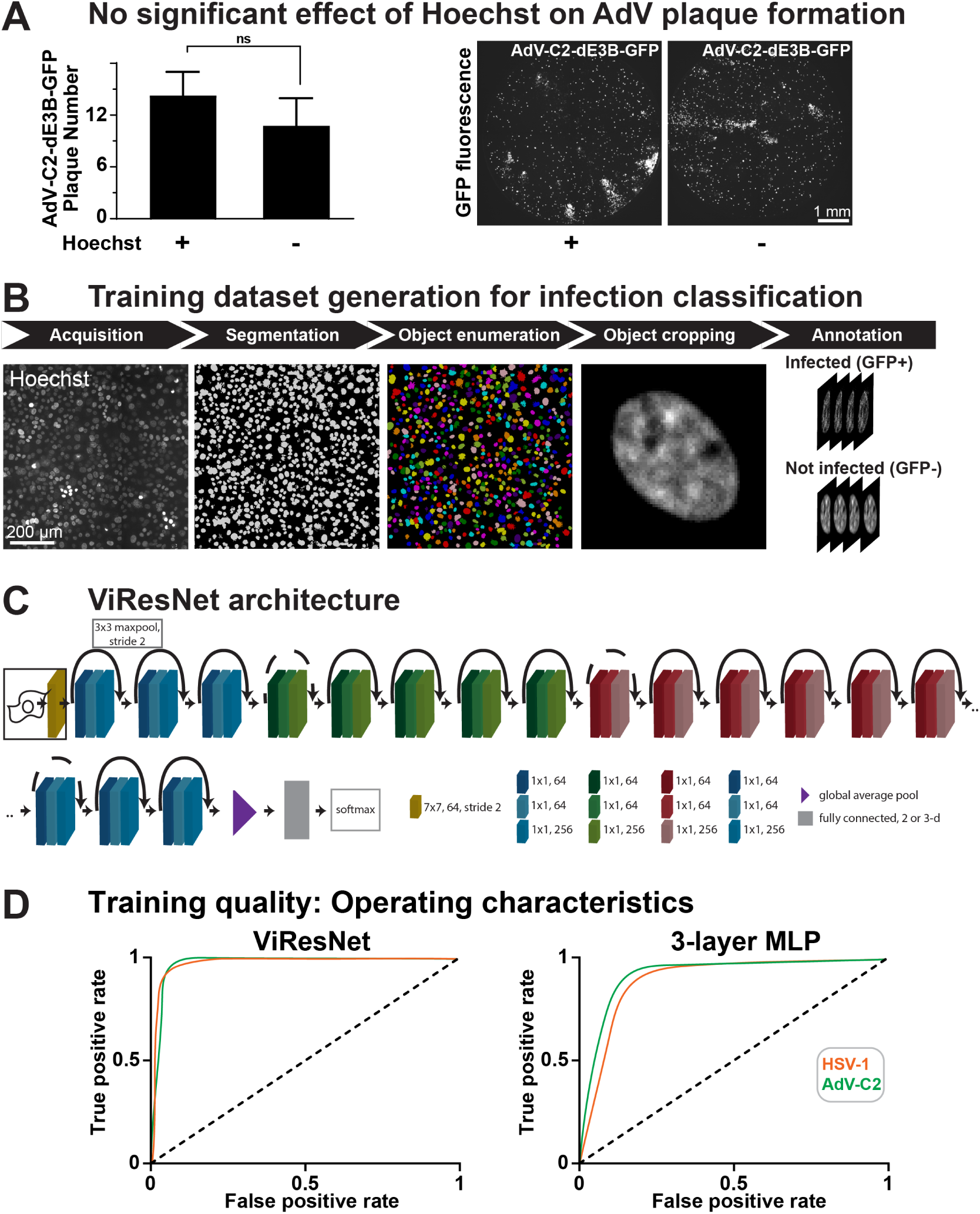
CNN architecture, dataset generation pipelines and training quality of ViResNet for infection classification. (related to Fig. 1). (A) Effect of Hoechst on AdV Plaque formation. (B) Training dataset generation pipeline for infection classification. (C) ViResNet architecture schematic. (D) ROC curves for infection classification (AdV, HSV-1) of trained ViResNet and multilayer perceptron.

**Sup. Fig. 2:**
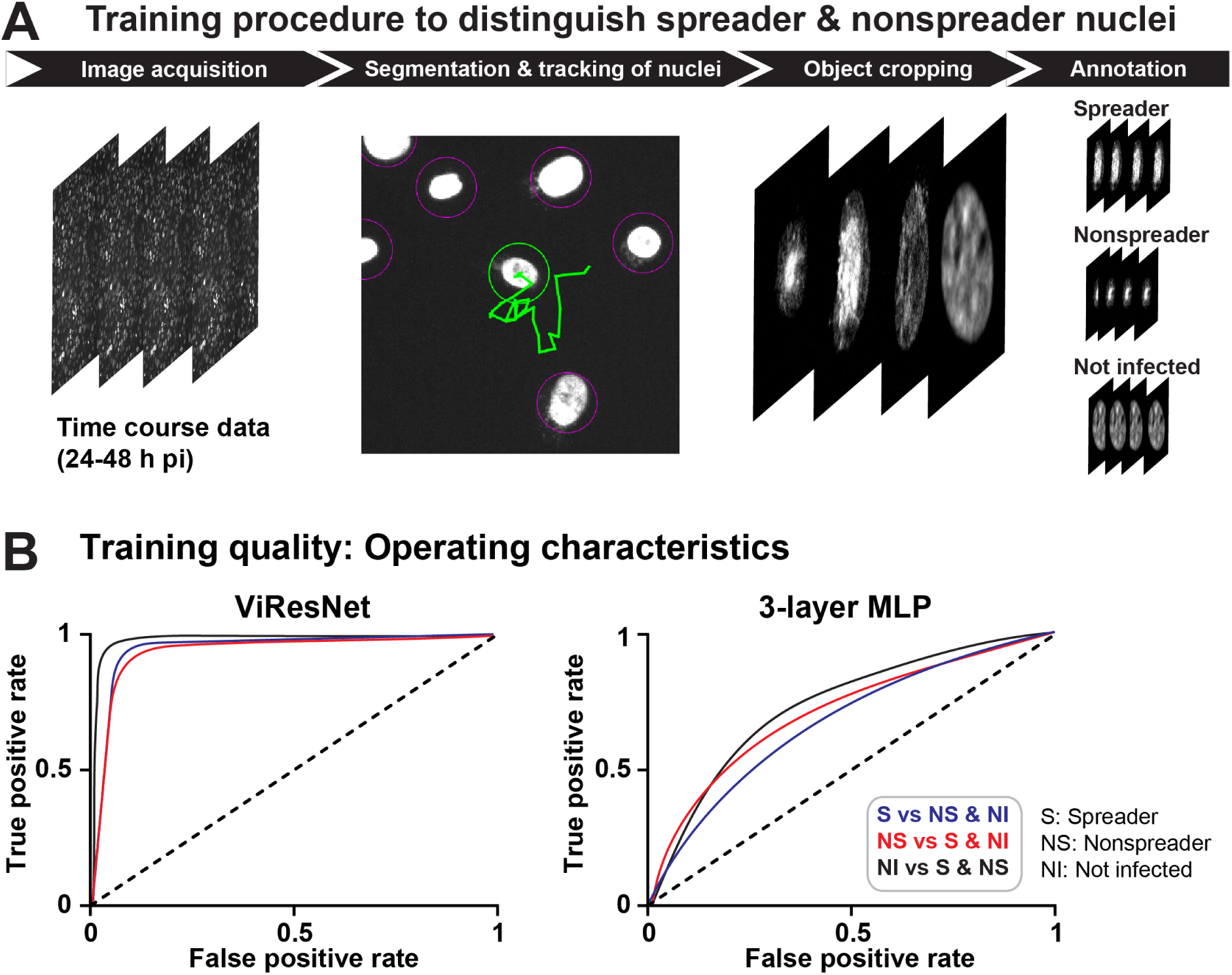
Dataset generation pipelines and training quality of ViResNet for infection prediction. (Related to Fig. 3). (A) Training dataset generation pipeline for prediction of spreading and nonspreading infected nuclei, respectively. (B) Receiver operating characteristic (ROC) curves for infection prediction of trained ViResNet and multilayer perceptron.

**Sup. Fig. 3:**
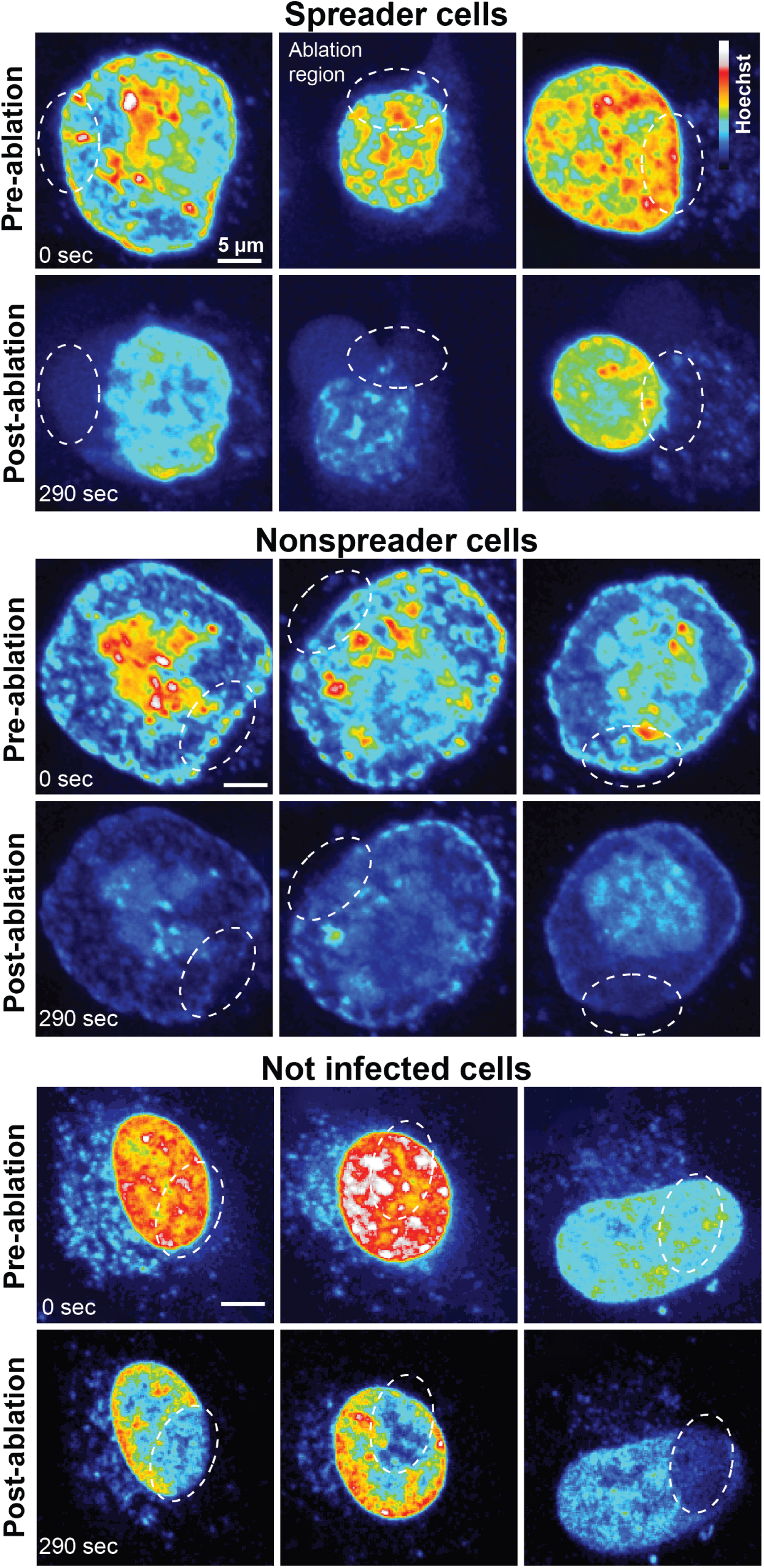
Samples of prospective spreader, nonspreader and not infected nuclei at pre- and post-ablation stages. (Related to Fig. 5). All images are represented as Z-stacks with Nyquist sampling.

***Supplementary Movies S1 - S3: Live cell analyses of AdV-C2-GFP-V or HSV-1-VP-16-GFP dynamics and lytic egress..*** (Related to Fig. 2 and Fig. 3).

HeLa cells stably expressing H2B-mCherry (47) were infected with AdV-C2-GFP-V at MOI 0.2 and recorded by live cell confocal fluorescence microscopy from 24 to 45 h pi at 15 min intervals. Merged pseudo-colored maximum projections are shown with GFP-V in green, and H2B-mCherry in red.

**Movie S1** Dispersion of GFP-V clusters from the nucleus into a confined cytoplasmic region around the nucleus (see cell in the center of the movie) 35:45 – 38:30 h:min pi. Note that the clusters and nuclear GFP-V signals disappear almost completely while the GFP-V signal in nonlytic infected cells persists. Related to Fig. 2.

**Movie S2** Loss of GFP-V clusters from AdV-C2-GFP-V infected cell nuclei. Related to Fig. 2.

**Movie S3** Nuclei are stained with Hoechst (blue), HSV-1-VP-16-GFP infection is visualized by the virus expressing VP-16-GFP fusion protein (green), and the nuclei of lysed cells are stained with PI (red). Related to Fig. 3.

